# Comprehensive collection and prediction of ABC transmembrane protein structures in the AI era of structural biology

**DOI:** 10.1101/2022.07.08.499254

**Authors:** Hedvig Tordai, Erzsébet Suhajda, Ian Sillitoe, Sreenath Nair, Mihály Váradi, Tamás Hegedűs

## Abstract

The number of unique transmembrane (TM) protein structures doubled in the last four years that can be attributed to the revolution of cryo-electron microscopy. In addition, AlphaFold2 (AF2) also provided a large number of predicted structures with high quality. However, if a specific protein family is the subject of a study, collecting the structures of the family members is highly challenging in spite of existing general and protein domain-specific databases. Here, we demonstrate this and assess the applicability and usability of automatic collection and presentation of protein structures via the ABC protein superfamily. Our pipeline identifies and classifies transmembrane ABC protein structures using PFAM search and also aims to determine their conformational states based on special geometric measures, conftors. Since the AlphaFold database contains structure predictions only for single polypeptide chains, we performed AF2-Multimer predictions for human ABC half transporters functioning as dimers. Our AF2 predictions warn of possibly ambiguous interpretation of some biochemical data regarding interaction partners and call for further experiments and experimental structure determination. We made our predicted ABC protein structures available through a web application, and we joined the 3D-Beacons Network to reach the broader scientific community through platforms such as PDBe-KB.

## Introduction

ABC (ATP Binding Cassette) proteins play crucial roles in diverse biological functions from bacteria to man [1,2]. Importantly, human ABC transmembrane proteins are involved in several pathophysiological processes [1,3,4]. Mutations can affect their folding, assembly, trafficking, plasma membrane stability, and function, leading to decreased functional expression [5,6]. Therefore, understanding the effect of mutations on their structure and dynamics is highly important. Several family members extrude xenobiotics (e.g. toxic molecules and therapeutic drugs) from the cell, thus, learning the substrate recognition mechanism of these multidrug transporters can improve drug development and may prevent unwanted drug interactions [2,7]. Consequently, determining the atomic level 3D structure of ABC proteins is a widely researched area [8–10]. For example, various steps of structure-based drug design, such as selecting the appropriate protein target, understanding its pathological mechanism at the molecular level, and developing a new therapeutic substance or redesigning a drug, requires the knowledge of the target protein structure at the atomic level [11].

The major protein structure determination method was X-ray crystallography for several decades [12]. This technique requires an enormous amount of resources, especially in the case of transmembrane proteins, which are the target of a large fraction of prescription drugs [13,14]. The examined protein needs to be isolated and purified in a large quantity and the process of crystallization requires a trial-and-error approach [15]. Furthermore, it is rare to obtain structural information on the membrane environment from the crystal. In recent years, cryo-electron microscopy (cryo-EM) emerged and became the ultimate structure determination method for transmembrane proteins [16]. The cryo-EM maps may also include information about the membrane environment [17]. Despite these recent advances in the structure determination of transmembrane proteins, they only make up ∼5% of all the protein structures in the Protein Data Bank, while close to 50% of prescription drugs target TM proteins [13,14].

Researchers utilized enormous resources in the last decades to predict protein structures from sequence information [18]. In recent years, an algorithm based on deep learning, AlphaFold2 emerged with remarkable accuracy in this task [19]. Its high quality and speed of structure prediction resulted in a vast database of predicted structures, the AlphaFold Protein Structure Database, which currently holds close to 1,000,000 protein structures [20]. AlphaFold2 (AF2) is serving now as an ultimate tool for learning protein structures which would be hard to resolve experimentally, like protein complexes and transmembrane (TM) proteins. Although AF2 was not trained to predict transmembrane proteins, we have recently demonstrated on ABC proteins that this novel deep learning method is able to provide high-quality structures of transmembrane proteins [21]. RoseTTAFold [22] and trRosetta [23] have also been published as high-accuracy structure predictors, based on various deep learning approaches.

ABC proteins are an excellent group of proteins to study AF2 performance on TM proteins [21] since their TM domains are not conserved because of their heterogeneous transport functions. Therefore, the structures of their transmembrane domains exhibit various scaffolds that currently enable transmembrane ABC structures to be clustered into nine distinct structural classes (Figure S1) [24]. In contrast, the nucleotide binding domains (NBDs) of ABC proteins are highly conserved, and contain Walker sequences and the ABC signature or fingerprint [8]. Notably, an ATP binding site is formed by the Walker A and B from one NBD and signature from a second NBD. Therefore, the functional form of a TM ABC protein includes two NBDs and TMDs. These domains can be encoded in one polypeptide chain in the case of full transporters, in two chains in the case of half transporters working in dimeric forms, or in four chains [25,26]. The structures of ABC proteins can be observed in various conformations [8,27]. For example, proteins with Pgp-like structures (Figure 1) exhibit a widely open conformation towards the intracellular space, capable of substrate binding, in the absence of ATP. When two ATP molecules are bound, the NBDs close and form a tight interaction and the extracellular ends of TM helices open to provide a dissociation site towards the extracellular space. These conformations are associated with the steps of the alternating access mechanism [9,28]. Other types of proposed mechanisms for ABC proteins include the peristaltic or credit card swipe mechanisms. For transporters operating with these mechanisms, the aforementioned two distinct conformations cannot be distinguished so promptly. [8,29,30].

**Figure 1:**
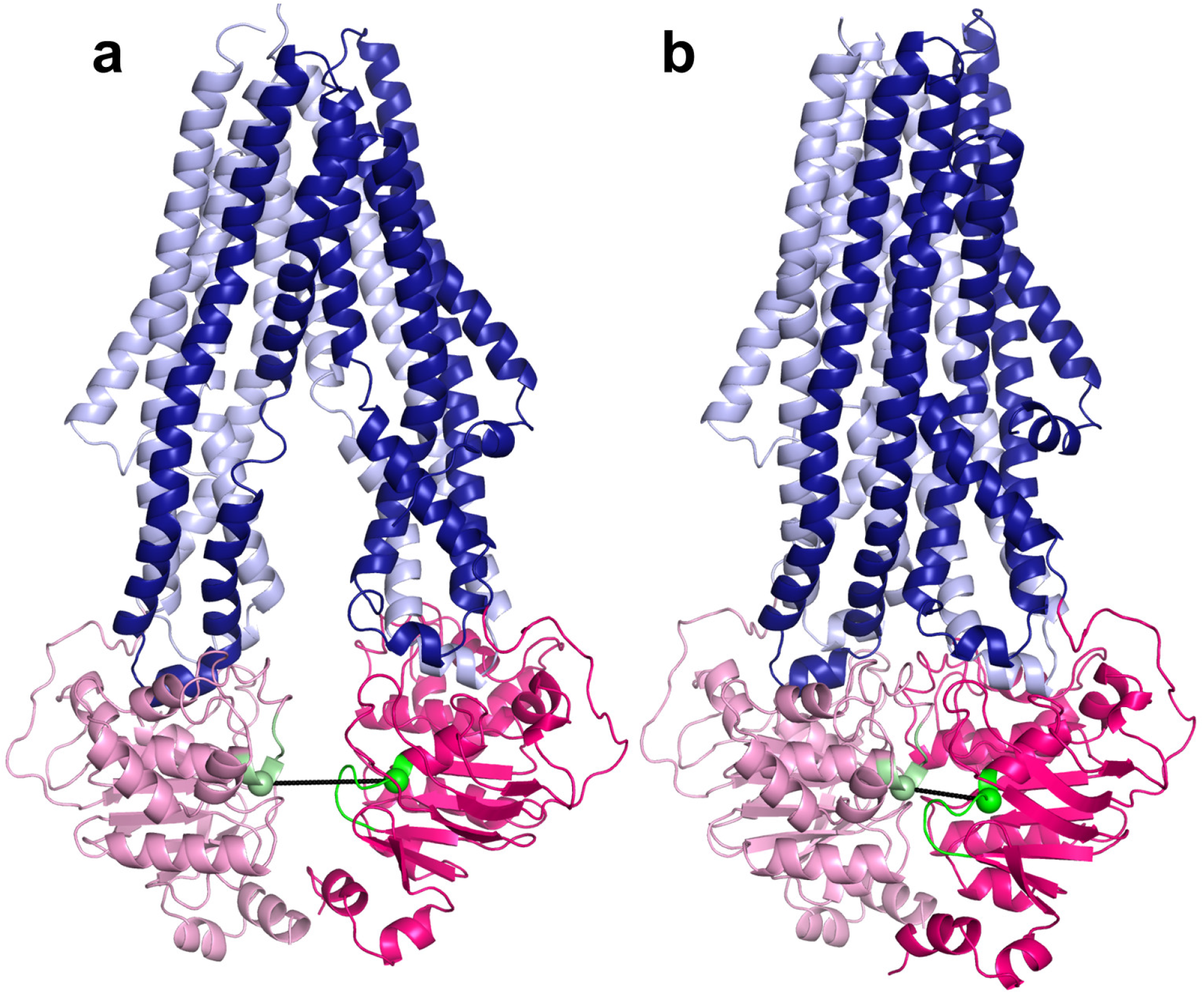
Inward-facing (a, PDB ID: 7a69) and inward-closed (b, PDB ID: 6c0v) structures of ABCB1/MDR1/Pgp. Light and dark blue colors: TM domains; pink and hot pink: NBDs; green and pale green: the last residue in the Walker A and ABC signature helices, respectively; black: the conftor/distance between the selected amino acids from Walker A and signature.

Although various structures of numerous ABC proteins were determined in the last years using cryo-electron microscopy, 25 of the 44 physiologically and pathologically important human ABC transmembrane proteins lack experimental structures in the PDB. The AlphaFold DB, co-developed by EMBL-EBI and DeepMind [20], supplemented the set of resolved transporters, albeit only full-length transporters, since only single-chain structures are available from this database. While the significant increase in the number of reliable structures for transmembrane proteins is a welcome change, the fact that the structures have to be collected from multiple data providers poses a potential challenge to performing comparative computational and experimental studies. Although both PDB and AF2 structures can be searched with a structure for homologous ones using the DALI server [31], it cannot reliably distinguish different ABC TM folds (https://abc3d.hegelab.org/dali.html). A different approach to solve this issue is the use of the SCOPe domain database [32], but it contains only three types of ABC folds (f.22: ABC transporter involved in vitamin B12 uptake, f.35: Multidrug efflux transporter AcrB transmembrane domain, and f.37: ABC transporter transmembrane region). To tackle the challenges associated with ABC protein structure collection, we built an automated pipeline that collects ABC protein structures from both the PDB and AlphaFold DB [20,33]. We supplemented the structures with relevant metadata (e.g. conformational state, structural family, method of resolving structure) and made them available through a web application (https://abc3d.hegelab.org). We also predicted the functional unit structures of human half transporter dimers using AlphaFold-Multimer [34]. In order to make our data set more accessible, we also linked our service (https://3dbeacon.hegelab.org) to the 3D-Beacons Network (https://3d-beacons.org).

## Methods

### Databases and associated software

Experimental structures and associated sequences (release of 2021-12-29) were downloaded from RCSB. AlphaFold predicted structures of 21 proteomes were retrieved from the AlphaFold Structure Database in July, 2021 [20]. These sets were used for analysis in this study, except for a few cases, which raised after updating our database with current experimental structures and the AF structures covering proteins in SwissProt.

ABC PFAM entries were identified at https://pfam.xfam.org (n=29) and extracted from the Pfam-A.hmm file. The selected entries and their accession numbers are listed in our previous publication [21] and can be downloaded from http://abc3d.hegelab.org/pub/abcdoms.hmm. The sequence of every structure in the PDB and AF datasets was searched using HMMER hmmsearch (http://hmmer.org) [35]. The E parameter was set to 0.001 and the match length was restricted to a minimum of 90% of the HMM profile length. The hmmsearch output was parsed using BioPython [48].

### Identification and classification of TM ABC proteins

Entries containing at least one TM Pfam hit and at least two NBD Pfam hits were selected and classified as transmembrane ABC proteins. Based on this comprehensive collection, corresponding biological unit files (.pdb1) and structural prediction files (.pdb) were downloaded from PDB and AlphaFold Structural Database, respectively. The downloaded structures were then classified into structural families based on both Pfam hits and structural similarity to selected reference structures, listed in Table S1. From the sequence search results, PDB entries that included at least two NBDs and one TMD in the biological unit file (encoded in one or more chains) were selected as functional ABC protein structures. Then, TM-align was performed aligning every single structure to the reference structures, using TM-score for characterizing structural similarity and to assure the reliability of the classification [37]. We considered two structures belonging to the same structural class if the TM-score resulted from their alignment was above 0.6. E.g. the Bce-like family emerged as a MacB Pfam hit with low structural similarity to the MacB reference structure (TM-score of 0.49).

To further classify the structures into inward-facing and inward-closed conformations, we used the distance between the Cα atom of the terminating amino acid of the Walker A helix (GXXXXG**<u>K</u>**T) in one of the NBDs and the signature helix (LS**<u>G</u>**GQ) in the other NBD. Since the signature sequence can be degenerate and cannot be recognized by sequence search, we identified these positions based on structural alignments with the Pgp NBD1 from 6c0v, using TM-align [37]. Structures were analyzed using MDAnalysis [38] and NumPy [39].

### A pipeline for automatic updates

In order to keep our collection updated, a Python script checks the sequence files deposited in PDBe and AlphaFold DB weekly. In case of changes, the aforementioned steps are executed automatically on the sequence of new entries. If the algorithm finds two NBD Pfam hits and no TMD hit in the sequences of the chains, we manually check the corresponding structure to avoid missing structures with novel transmembrane folds. Additional data needed for classification and the determination of conformational states is collected from cif files and UniProt [40]. The collected information is stored in a PostgreSQL database (https://www.postgresql.org). To make the appearance of structures on our website more uniform, the pdb files of identified ABC structures are aligned to the corresponding reference structure, regarding both structural family and conformational state, using the rmsd function of PyMOL (The PyMOL Molecular Graphics System, Version 2.4.0 Schrödinger, LLC). The final pdb files were converted to mmCif files with gemmi [41].

### Web applications

Both the 3dbeacon and the abc3d web applications are based on FastAPI (https://fastapi.tiangolo.com) placed behind an nginx (https://www.nginx.com) web server. All data for our 3dbeacon client is stored in json files. The abc3d application data are tied to the PostgreSQL database via the SQLAlchemy object-relational mapper [42]. Web layout strongly depends on JavaScript, JQuery (https://jquery.com), and bootstrap (https://getbootstrap.com).

### Running AlphaFold2

AlphaFold initial release and v2.0.0 were downloaded from github and installed as described (https://github.com/deepmind/alphafold) under Linux (Debian 10, 96GB RAM, NVidia Quadro P6000 GPU with 24 GB RAM or NVidia RTX A6000 GPU with 48GB RAM GPU). We introduced minor modifications into the code to overcome memory usage problems in case of large multiple sequence alignment files and to be able to run multimer predictions with the initial release (http://alphafold.hegelab.org). Our runs used all genetic databases (*--db_preset=full_dbs*). Generated structures were evaluated based on PAE, pLDDT, and ipTM+pTM scores [19,34]. In addition, all top scored structures were inspected visually.

### Data visualization

Molecular visualization was performed using PyMOL (The PyMOL Molecular Graphics System, Version 2.4.0 Schrödinger, LLC). Graphs were generated using Python’s *matplotlib* library [43].

## Results and discussion

### Connections between transmembrane ABC proteins structures and Pfam profiles

Running our pipeline yielded 325 PDB structures (as of 29/12/2021). We selected the top scoring TMD matches of our Pfam searches as the base for classification (Figure 2). We found only three incorrect biological unit files (3wme, 3wmf, and 3wmg), which contained only half of the functional unit of an ABC transporter (1 NBD and 1 TMD). Instead of numbered structural classes [24], we use names related to the most known representative member of a class (e.g. Pgp-like and BtuCD-like) (Figure S1), since we find this not only more informative than numbers, but the recently renumbered classes cause confusions in publications of the last decades (e.g. type I ABC exporter in the old system corresponds to type IV transporter structure in the recently proposed system [24]). In addition, the numbering strongly suggests an order of the structural families (e.g. an ordering based on the evolutionary level of TM domains) but that is not the case.

**Figure 2:**
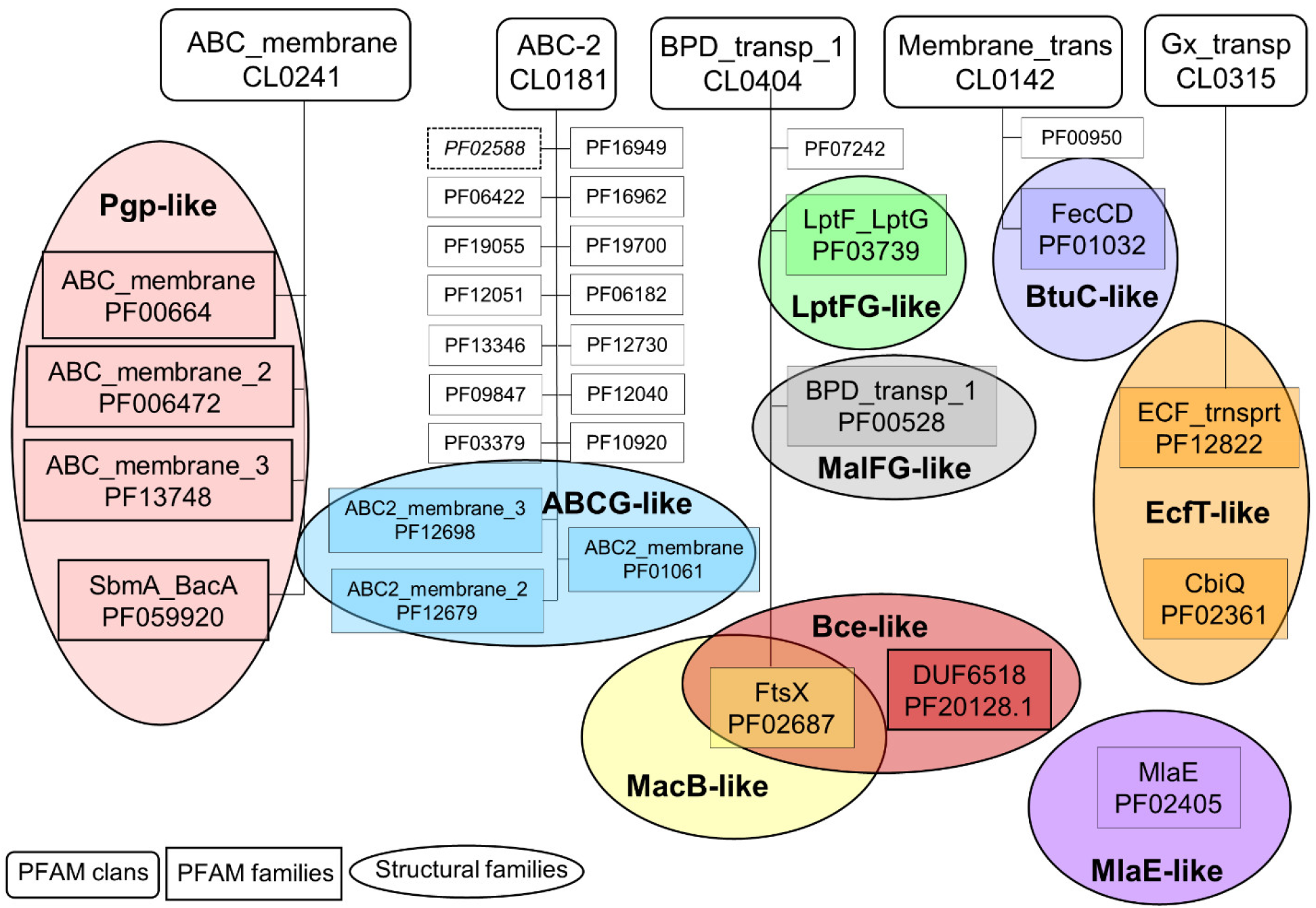
Pfam and structural classes of ABC proteins. Pfam clans (rounded boxes) with ABC protein hits are depicted. Those Pfam families (boxes), which include proteins with experimental structures are circled, colored, and labeled in bold. The labels were selected based on a widely known member of the structural family (e.g. Pgp-like). PF02588/YitT_membrane is in italic and with dotted outline, since it likely does not involve ABC family members.

The investigation of sequence and structure conservation led to important and interesting information regarding this diverse protein family. The HMM search showed matches in five Pfam clans (Fig.2.). The *ABC_membrane* clan includes four PFAM families, exhibiting Pgp-like structures. Of these four families, most of the existing experimental structures belong to the *ABC_membrane* family. *ABC_membrane2* family contains ABCD [44,45] and a mycobacterial ABC transporter, Rv1819c [46] structures, while no experimental structure is associated with the *ABC_membrane3* family. However, trRosetta and AlphaFold2 predictions also suggested Pgp-like architecture for members of this family (e.g. https://alphafold.ebi.ac.uk/entry/Q5F7D3). Importantly, we found that automatic computational models need careful interpretation. The trRosetta prediction of a protein of this Pfam family (UniProt ACC: Q9CNM3) did not contain the full sequence. In addition, neither the trRosetta nor the AlphaFold prediction took the dimeric nature of this protein into account (Figure S2). Interestingly, a member of the fourth family (*SbmA_BacA*) was demonstrated to form a homodimer of two TMDs and to function without any NBDs, thus it cannot be considered a real ABC protein (PDB ID: 7p34) [47].

The *ABC-2* clan includes proteins which exhibit, or potentially exhibit ABCG-like structures. Most of the related experimental structures belong to the *ABC2_membrane* and *ABC2_membrane_3* families including ABCG and ABCA proteins, respectively. The *ABC2_membrane* Pfam entry matches the first five TM helices of ABCG proteins. The exclusion of TM6 is most likely caused by the non-conserved extracellular loop between TM5 and TM6. TM1 was also not matched in most ABCA TMD by the *ABC2_membrane_3* profile, which involves a portion of their large extracellular loop (Figure S3). Experimental ABCG-like structures (6xjh and 6xji) in the *ABC2_membrane_2* family were determined for the same protein, PmtCD, exporting bacterial toxins [47]. The remaining families involve trRosetta or AlphaFold2 computational models. Most of the sequences of these proteins incorporate transmembrane domains and some other domains (e.g. Sodium/calcium exchanger, AAA, Dihydrodipicolinate synthetase, and Phosphotransferase enzyme in the CcmB and DUF3533 Pfam families), which are not characteristic for ABC proteins. These proteins may or may not function without NBDs and are good targets for studying the evolution of ABC proteins. In contrast to other *ABC-2* clan members, the Pfam profile of ABC2_membrane_7, PDR_CDR, and YitT_membrane (PDB ID: 3hlu) does not match membrane domains, but soluble regions instead. In addition, the AF2-predicted TMD of YitT (e.g. https://alphafold.ebi.ac.uk/entry/O34792) does not resemble an ABCG-like structure, thus this family likely does not belong to this group of proteins [21].

The *BPD_transp_1* clan includes four families with diverse structures (Figure 2, Figure S1). Three Pfam families include proteins with experimental structures belonging to different structural classes (MacB-, Bce-, LptFG-, and MalFG-like). Notably, the known Bce-like structures (PDB IDs: 7tcg and 7tch [48]) involve two transmembrane domains with MacB-like and Bce-like folds on a single polypeptide chain. The fourth family, DUF1430, is defined by a soluble domain. In these proteins, cytosolic domains likely prohibit the binding of NBDs to the TMDs, thus they may not be considered real ABC proteins.

Members of both the *ECF_trnsprt* family (*Gx_transp* clan) and the *CbiQ family* (no parent clan) exhibit an EcfT-like structure. The *FecCD* and ABC-3 families in the *Membrane_trans* clan possess BtuCD-like structures.

### Conformational states of ABC protein structures

Since for ABC proteins two major conformational states can be distinguished [9,28], we also aimed to group the structures according to their inward-facing or outward-facing state. Since the latter state is related to ATP-induced association of the two NBDs, theoretically we could divide the structures into these two conformational states based on the presence or absence of nucleotides in the structure. However, there are several structures with bound nucleotides (e.g. 6z5u, 7oj8, 7ojh, and 7oz1), exhibiting separated NBDs, caused most probably by experimental conditions (Figure S4). For example, in the 6z5u structure, the non-hydrolyzable ATP analogue, AppNHpANP was not sufficient to trigger the transition from bottom-open to bottom-closed state [49]. As another example, a recent ABCG2 structure was determined under turnover conditions and the presence of ATP did not drive a complete closure of the two NBDs (Figure S4) [50].

Since the presence or absence of nucleotides proved to be an inefficient way of categorizing structures, we then tried to do so by calculating the RMSD between each individual structure and two typical reference structures representing either inward- or outward-facing conformations of the structural family the protein belongs to (Table S1). All the structures and their labels were manually validated after classification. We found higher number of mislabeled structures only in the Pgp-like family, where 18 out of 118 inward-facing structures were classified as outward-facing. These RMSD calculations also uncovered some flaws in currently used structural alignment tools demonstrated in Figure S5.

We have developed 3D vector based measures, conftors, to describe the relative orientation of domains and highlight structural differences in ABC protein structures [51]. Since the inward- and outward-facing conformation correlates with the level of NBD association (Figure 1 and Figure S1), we defined |conftor(WA/SIG)|, the distance between the Walker A in one of the NBDs and the signature motif from the other NBD to classify the conformations (Figure 1). For this definition, the Cα atom of the terminating amino acid of the Walker A (GXXXXG**<u>K</u>**T) and the signature (LS**<u>G</u>**GQ) helices were selected. We did not select the amino acids participating directly in ATP binding, since they are in loops, potentially causing noisy fluctuations in distance values. These positions were identified by aligning the query NBDs to a reference NBD (Pgp NBD1 from 6c0v) using the TM-align algorithm and not by simple sequence search, since there are many ABC proteins with degenerate, non-conserved ATP-binding sites (e.g. CFTR/ABCC7 possesses a degenerate signature motif [52,53]). The distribution of the distances between Walker A and signature calculated for all structures (Figure 3) indicated a cutoff value of 14 Å for distinguishing bottom-closed and bottom-open conformations. Indeed, visual inspection of structures with |conftor(WA/SIG)| = 14 ± 2 Å confirmed this value as a rational cutoff. We aimed to identify the source of very large NBD separation, thus we compared the distance values of X-ray and cryo-EM structures, since the large separation may have been the result of non-physiological crystal contacts in X-ray structures. We observed distances larger than 51 Å only in structures determined by X-ray crystallography (Figure S6).

**Figure 3:**
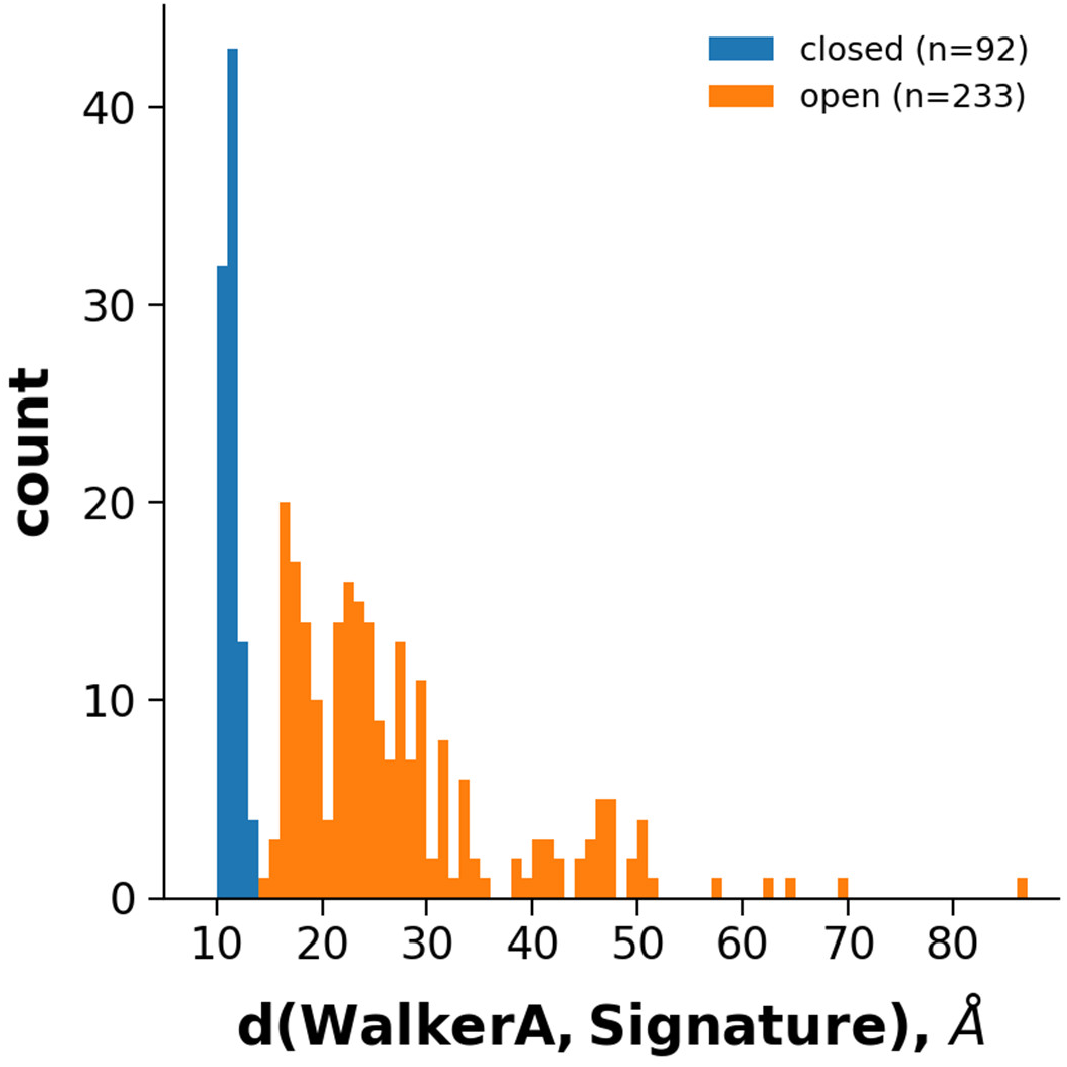
Grouping experimental structures based on the length of conftor(WA/SIG). The distribution of distance values of all collected ABC protein structures.

This type of structural classification to inward-facing and outward-facing categories is important, since the opening level and the dynamic alterations in the opening affect the substrate access to the central binding pocket [54]. This is clearly evident in the case of some Pgp-like structures, which exhibit an ATP-bound conformation with a well-defined, large, outward open cavity (Figure S7) [55]. In contrast, the difference in the TMDs of ABCG structures is subtle between the apo and ATP-bound states (Figure S7). Although the TM helices are reorganized upon ATP binding, no large opening on their extracellular end is observed. It is also important to note that the static ABC structures cover different parts of a continuous space of possible conformations. This is highly prominent in the case of inward-facing conformations, which exhibit a wide range of opening levels (between 14 and 86 Å) and also various conformational states of the two NBDs relative to each other. We highlight two examples associated with this issue. First, ABCG2 (PDB ID: 6vxf) has an additional conformation named inward-facing-closed [56]. In this ATP-bound conformation the NBDs are separated, but the intracellular ends of some TM helices are somewhat more closed than in the inward-facing conformation (Figure S7). Although the difference between TM helix ends is subtle, it makes the large drug binding cavity in the TM region inaccessible from the intracellular space. However, since the overall conformation is almost identical to the inward-facing ABCG2 structures (RMSD = 1.736 Å) we classified this structure as inward-facing. Second, the NBDs in EcfT proteins show a closed conformation both in the ATP-bound (5d3m) and apo structures (4huq, 4rfs, 4hzu, 5jsz, 5×3x, 5×41, 6fnp, 6zg3, 7nnt, and 7nnu), which is most likely caused by their special TMDs and an unconventional transport mechanism [57,58]. Namely, in the absence of a substrate, one of the TM domains, the substrate binding S-factor, is not associated with the complex consisting of the EcfT TM domain and the two NBDs, thus EcfS cannot keep one of the NBDs in the complex. Therefore, other interactions are needed, including a mechanism keeping the NBDs in contact in the absence of ATP as well. Most of the WA/SIG distance values (19 out of 2 × 11) of EcfT transporters are between 14 and 19 Å and three are slightly below 14 Å.

### ABC protein structures predicted by AlphaFold2

The AlphaFold Protein Structure Database contains predicted structures for most of the protein sequences in the UniProt database. We collected ABC protein structures from this AF database following the same procedure as in the case of the experimentally determined structures from the PDB, namely by using Pfam searches. However, since the AlphaFold DB only contains monomeric structures, we gathered only full transporters, which encoded two NBDs and two TMDs within a single polypeptide chain. As a result, the hits belonged only to Pgp-like and ABCG-like structural classes. While the open structures were overrepresented in the AF-prediction of 21 proteomes (Fig. 4), the numbers of open/closed conformations are more balanced in the prediction of the SwissProt dataset (101 vs. 414 and 163 vs. 136 in the Pgp-like and ABCG2-like families).

**Figure 4:**
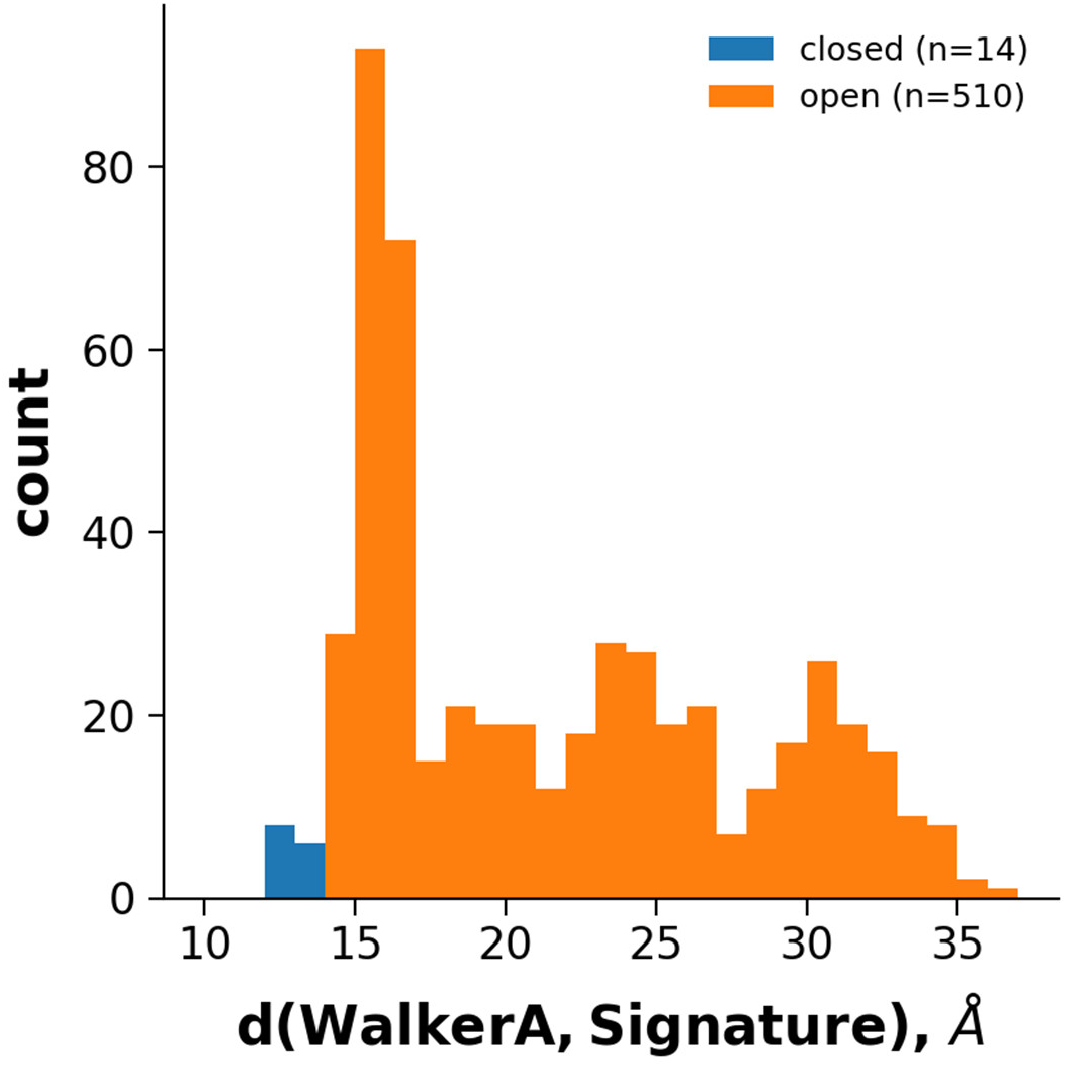
Grouping AF2 structures based on the length of conftor(WA/SIG). The distribution of distance values of all collected AF2-predicted, full ABC structures.

We noticed that there is no available experimental or AlphaFold-predicted structural data for some human dimeric ABC proteins. Since we found it critical to generate the functional forms of human ABC protein structures that can facilitate both learning their structure/function relationship and drug development, we predicted dimeric human ABC structures. Previously, we have shown that AlphaFold2 provided high-quality structures for both homodimeric and heterodimeric ABC proteins [21]. However, AlphaFold-Multimer has been recently released, and it has a significantly better performance on heterodimeric complexes when compared to the original AF2 [34]. Using AF-Multimer, we performed test multimer runs for ABC dimers with known structures (e.g. ABCG5/ABCG8, PDB IDs: 5do7, 7jr7, 7r87-7r8b) that yielded high-quality structures (Table S2). Then we performed predictions for proteins with no available structures in the PDB or which had longer unresolved regions (Table S2). For example, several ABCB family members possess an additional N-terminal TM domain (TMD0) which are not resolved or exhibit low resolution (Figure S8). Interestingly, in our AlphaFold predictions there were no contacts between the core and TMD0, but the additional TMD0 helices were rationally oriented according to a putative membrane bilayer.

One of the most exciting tasks was to test if AF-Multimer was able to build the mitoSUR complex. Sulfonylurea receptors (SUR1/ABCC8 and SUR2/ABCC9) are octamers of four SURs and four inward-rectifying potassium Kir channels (Figure 5) [59,60]. Similar to ABCC8 and ABCC9, ABCB8 was found to regulate potassium current [61]. ABCB8 functions in the inner membrane of mitochondria and mitoK (alternative name: Coiled-coil domain-containing protein 51, CCDC51) was identified as its potassium channel partner [61]. In order to build the complex structure, we submitted eight copies of the half transporter ABCB8 and four copies of mitoK sequences to AF-Multimer. No reasonable structures were built. We also predicted the structure of four mitoK proteins without ABCB8. A complex was built, in which the pore was delineated by coiled-coil regions and the putative transmembrane helices were not aligned to match a lipid bilayer (Figure 5). Thus, the overall structure did not meet expectations of how a potassium channel should look like. Known potassium channels, even with low sequence similarity, exhibit two TM helices and a reentrant loop between these TM regions, which loop provides the selectivity filter with a characteristic amino acid pattern (TMxTVGYG) [62,63]. We could not find any similar potassium selectivity pattern in mitoK using less stringent regular expression patterns. There are two possible explanations for this phenomenon. On one hand, mitoK may be an additional regulatory factor for the potassium current regulated by ABCB8 and not the potassium channel itself. This seems unlikely, since the ABCB8/mitoK purified and reconstituted from bacteria exhibited potassium currents [61], and it is difficult to imagine this complex carrying a third protein (a potassium channel) during that complicated process. On the other hand, mitoK may represent a novel type of potassium channel, but AF-Multimer was not able to produce a meaningful prediction, since no previously known structure resembled this fold in its learning set. This is also unlikely in the light of our earlier study, demonstrating that AF2 was able to predict novel transmembrane folds not included in the learning and template sets [21].

**Figure 5:**
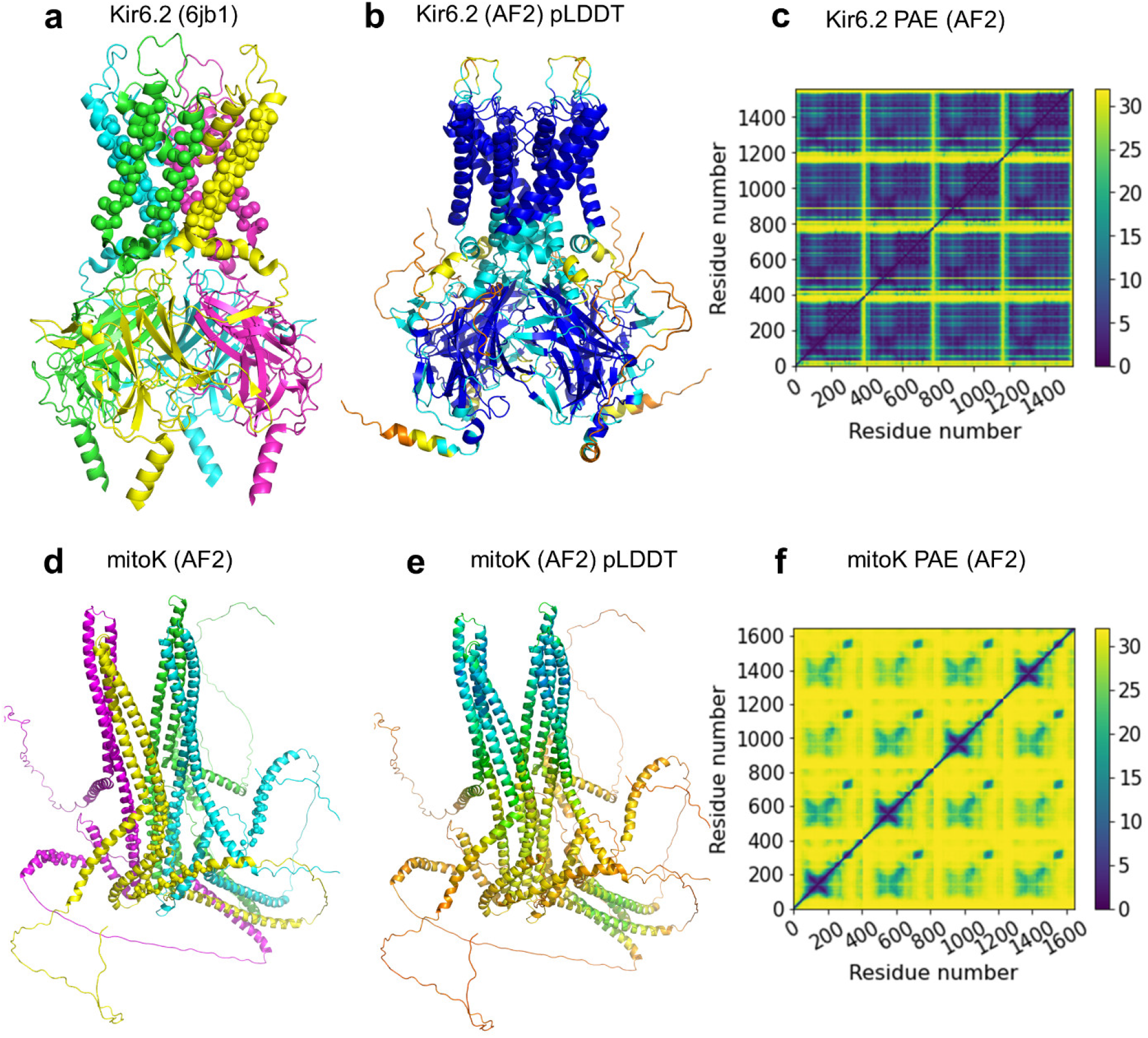
AF2 prediction of the mitoK tetramer. (**a**) The central tetramer of Kir6.2 potassium channel from the octamer structure with SUR1 (PDB ID: 6jb1). The four chains are colored differently and spheres indicate transmembrane regions. (**b**) The Kir6.2 tetramer predicted by AF-Multimer exhibit high pLDDT scores (blue and turquois regions). The RMSD calculated for TMDs was 0.8 Å when compared to its experimental structure. (**c**) The plot of predicted align errors (PAE) indicate low values thus a predicted structure with high quality. (**d**) The AF-predicted structure of mitoK, colored by chains, does not resemble a potassium channel architecture. Spheres indicate transmembrane regions. (**e**) The yellow and orange colors of the same structures correspond to poorly predicted regions with low pLDDT scores. (**f**) The high values of the PAE plot also indicate unreliable structure prediction for the mitoK tetramer.

### The ABC3D database

Since it is not trivial to access a list or a sublist of ABC proteins neither from the PDB nor from the AlphaFold Structural Database, we created a web application that contains the currently available full-length ABC structures (https://abc3d.hegelab.org). In this web application, structures can be browsed and searched in a categorized manner at various levels. On the first level, users can choose to display “Experimental”, “Computational” or “Human” proteins and they can also view a comprehensive list of every available 3D structure under the “All” menu item. Every category except Human is further grouped by structural classes (Pgp-like, ABCG2-like, MalFG-like, EcfT-like, BtuC-like, LptFG-like, MacB-like, Bce-like, and MlaE-like) and conformations (open or closed). In the Human category, we chose to group proteins by subfamilies (ABCA, ABCB, ABCC, ABCG, and ABCD) instead of structural classes. Not only is this arrangement more common in the field, but also every human ABC protein can be classified into just two structural classes: ABCA and ABCG proteins belong to the ABCG2-like conformations, while ABCB, ABCC and ABCD families exhibit Pgp-like structures.

One structural class or human subfamily can be selected by a single click, as well as structures with a specific conformation. Using the main Select button, more complex searches can be performed. Selections can be narrowed by not only structural class and conformation, but also by taxonomy, gene and protein names, and release dates. Each structure can also be selected or unselected individually. 3D structures can be visualized in the browser using Mol* [64] that helps the selection process. All structures are aligned to their corresponding reference structure that helps comparison.

Selected structures and associated data can be downloaded as pdb files and a tab -separated tsv format, respectively. mmCif files are also available for bulk download. All of the files are packed into a zipped file. Users can also opt to download only the metadata file. The main Download menu makes getting pre-assembled files for each structural class possible.

### Novel ABC structures accessible through the 3D-Beacons Network

In order to increase the visibility of our AF2-predicted ABC structures, we expose them through the 3D-Beacon Network (https://3d-beacons.org). This network provides access to both theoretical and experimentally determined structures. In addition to providing programmatic access to our models, we also have web pages for each individual model (e.g. https://3dbeacon.hegelab.org/uniprot/Q9H222/hege-abc-0001). Our implementation of the 3D-Beacons Client has some additional fields which are not present in the current standard definitions (https://3dbeacons.docs.apiary.io/#), so that users can have access to the methodology, publication, and coordinate files other than the main mmCIF file. We found the latter important, since some AlphaFold2 predictions contain structural regions which were most likely incorrectly built (e.g. soluble regions that enter the volume of the putative membrane region). In these cases, we published the trimmed version of the AF2-model as the main entry, but the full structure is also included on the webpage (e.g. ABCB10 homodimer, https://3dbeacon.hegelab.org/uniprot/Q9NRK6/hege-abc-0009). We also provide links to zip files containing all the structure files to download in different versions and formats easily.

## Conclusions

With the growing number of available structures in 3D databases, there is an increasing need for simple accession of several or all structures of a specific protein family. We developed a web application providing this functionality for transmembrane ABC proteins and our approach can serve as a prototype for accessing bulk structural data per protein families. Based on user feedback, it will be improved and standardized to build a general framework for accessing the 3D structural data of any protein family. The focus of the further developments will be on the high level automation of various steps (e.g. classification, structure collection, alignment). In this study we automatized the categorization of open/closed structures that was based on our manual curation of 325 experimental structures. In addition to existing ABC structures, we exposed novel, AlphaFold-generated, dimeric structures of half ABC transporters through an implementation of the 3D-Beacons Client (https://3dbeacon.hegelab.org). Besides current standard data fields, we found it important to attach extra information, such as methods and references to publications, which types of data are under heavy standardization by the 3D-Beacon community, extending the current fields.

Our results also highlighted some issues associated with the increasing number of available structures of proteins and protein complexes. First, characteristic conserved regions and structures should be identified more precisely than current approaches allow. Deep learning methods may allow domain identification with a higher accuracy than HMM profile search [65]. Second, high performance methods with novel and automatic logical processes are required to analyze and compare the structures of protein complexes efficiently (Figure S5). Our structure predictions strengthen our previous results [21] that AF models of TM proteins can be as high quality as that of soluble proteins. However, careful investigation of individual structures is required to help assess model quality and also to detect hints for interesting and important biological questions, such as the case of the mitoSUR/mitoK complex (Figure 5). Importantly, we made the set of human ABC protein structures complete with AlphaFold predictions that opens the way to merge and improve our ABC mutation database (http://abcm2.hegelab.org) [66,67] with this complete set of structures, thus interpreting any mutations of human ABC proteins in the context of their sequence and structure.

## Acknowledgments

This work was supported by NRDIO/NKFIH: K127961 and K137610, Cystic Fibrosis Foundation (CFF, HEGEDU20I0), NKFIH 2020-2.1.1-ED-2021-00179, and the Wigner Scientific Computing Laboratory (WSCLAB, the former Wigner GPU Laboratory).

