## Supplementary Material for "Comprehensive collection and prediction of ABC transmembrane protein structures in the AI era of structural biology"

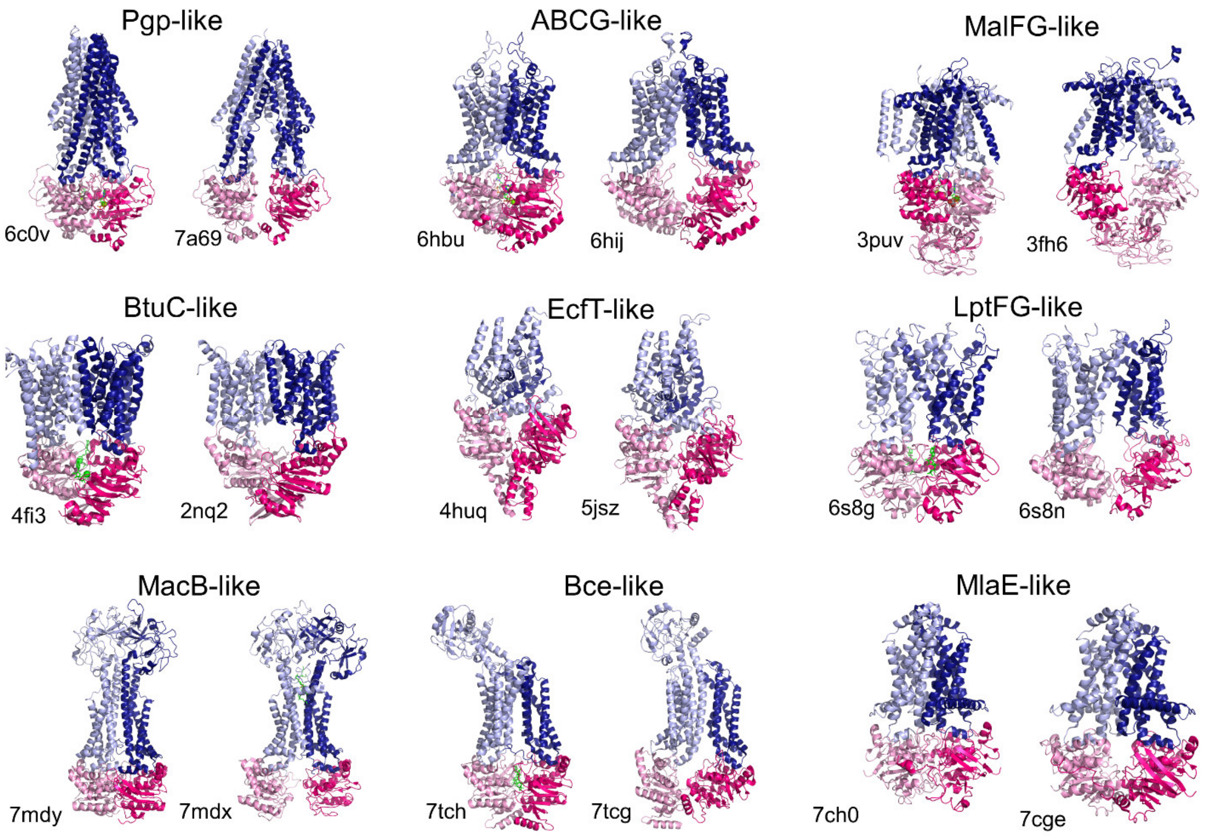


**Supplementary Figure S1: Currently known structural classes of ABC proteins.** A closed (left) and an open (right) structure are shown for each structural class. Pink and hot pink: TM domains; light blue and blue: NBDs; green: nucleotides.

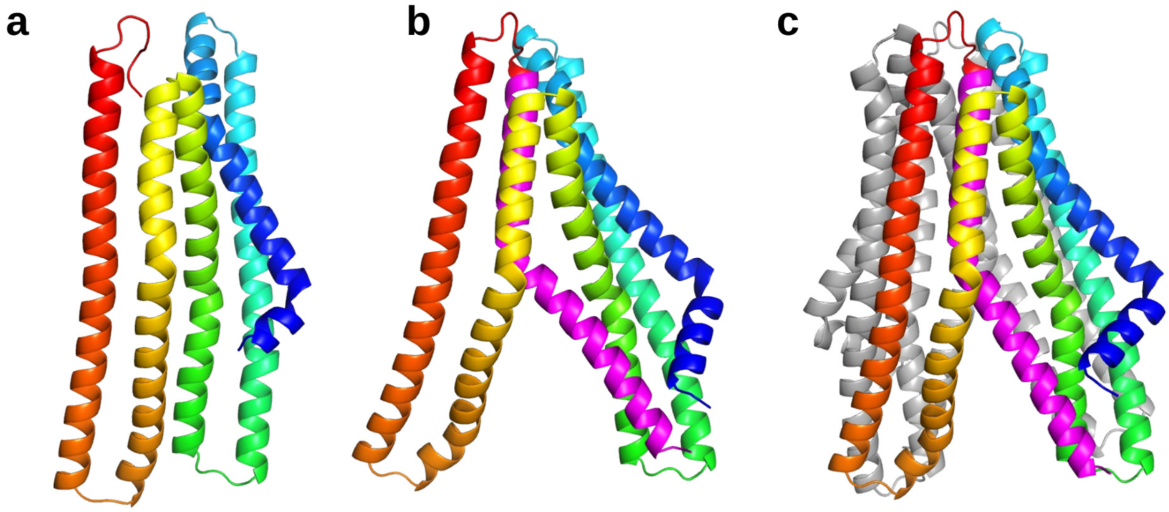

**Supplementary Figure S2: AI-predicted *ABC_membrane3* family (ABC_membrane clan) structures.** The structure of the TMD of an ABC protein from *Neisseria gonorrhoeae* (UniProt ACC: Q5F7D3) was predicted by trRosetta (**a**), AlphaFold2 (**b**), and Alphafold-Multimer (**c**). trRosetta prediction included only a.a. 7-243 corresponding to TM1 to TM5, colored from N- (blue) to C-terminus (red). Our AF2 prediction included the full length sequence (a.a. 1-288). The TM1 to TM5 region was colored from blue to red and TM6 was set to magenta. Gray in panel c: the second protomer. Interestingly, the AF2 monomer and the protomer in the dimer prediction are highly similar (RMSD = 1.75 Å).

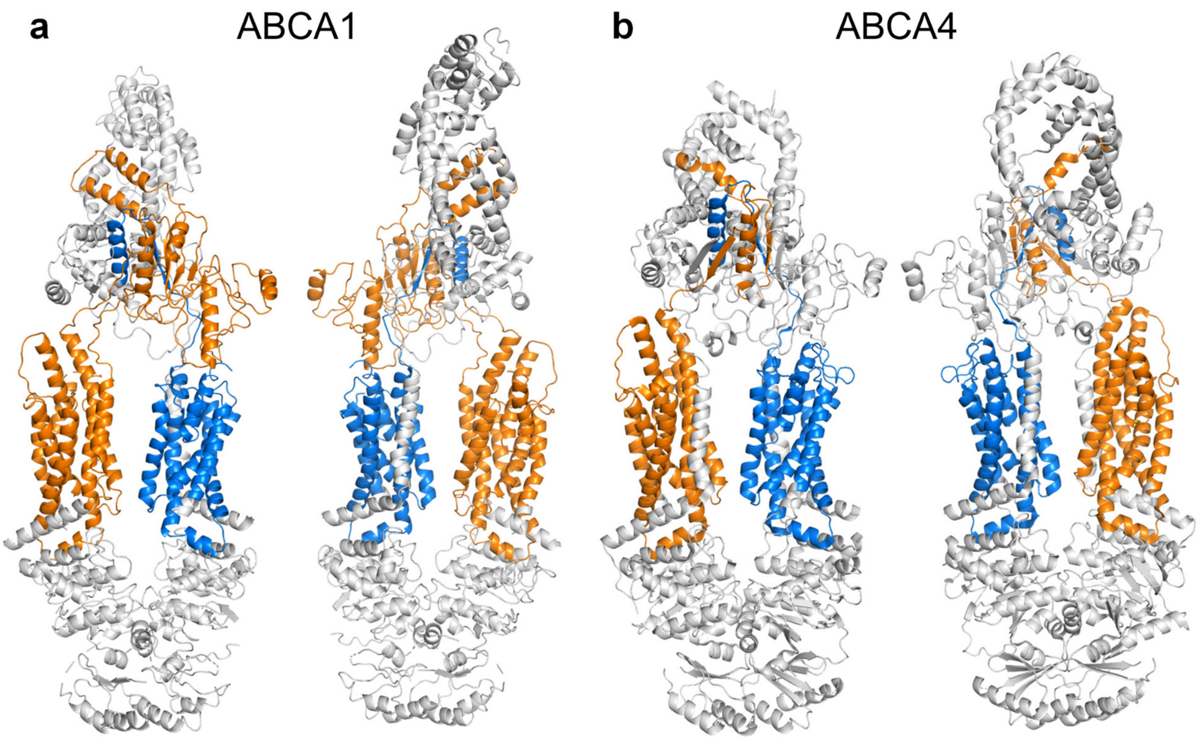


**Supplementary Figure S3: Pfam profiles may not match a complete domain.** (**a**) The N-terminal Pfam match did not include TM1 in ABCA1, but the C-terminal match covered TM7. (**b**) In ABC4 neither TM1 or TM7 were included in the PFAM matches. Blue and orange: N- and C-terminal Pfam matches, respectively. The Pfam profile PF12698 (ABC2_membrane_3) includes large extracellular regions between the first and second helix of both TMD. However, a region of the N-terminal extracellular loop is not matched, thus the first helix was not included in the match. This observation indicates that extracellular loops with low sequence conservation, which is common for loop regions, may interfere with automatic, accurate detection of domain boundaries using HMM profiles.

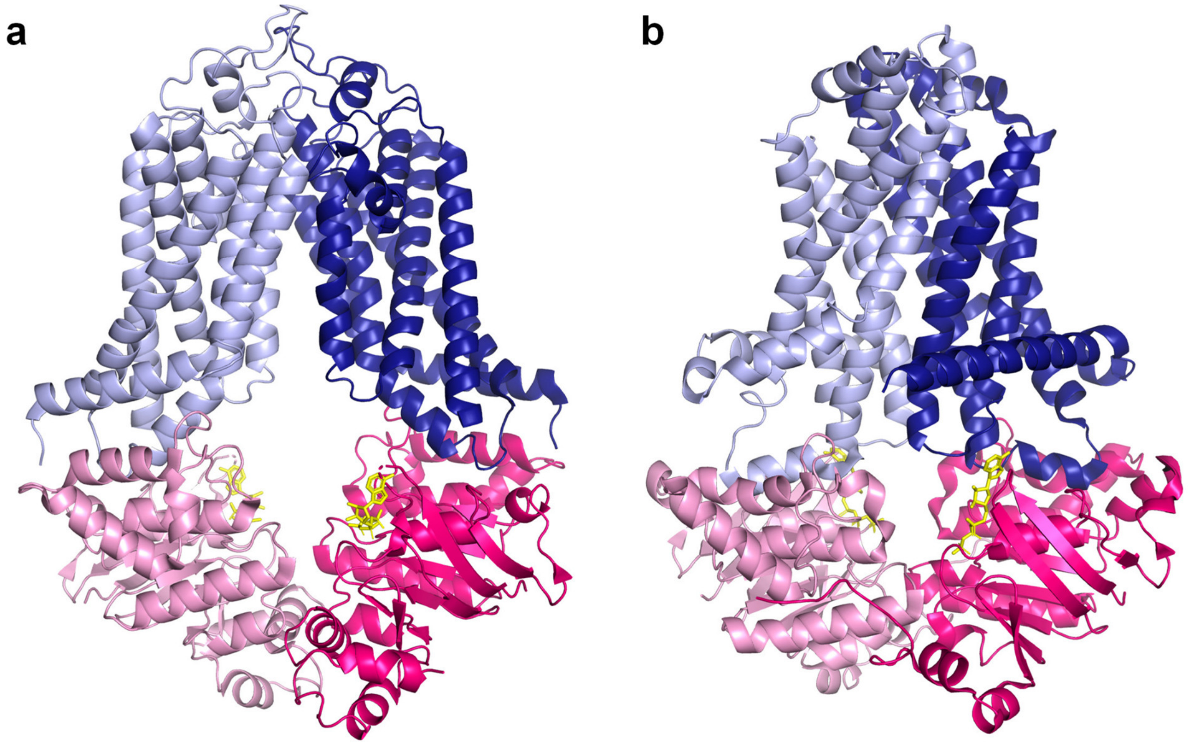


**Supplementary Figure S4: Inward-facing structures with bound nucleotides.** Several structures, determined in the presence of nucleotides, exhibit an inward-facing conformation. ABCG1 (**a**) (PDBID: 7oz1) and MlaBDEF (**b**) (PDBID: 6z5u) are shown as examples. In these cases, experimental conditions may not be sufficient to trigger the transition from inward-facing to the inward-closed state (34188171). Blue colors: TMDs, pink colors: NBDs, yellow: nucleotides.

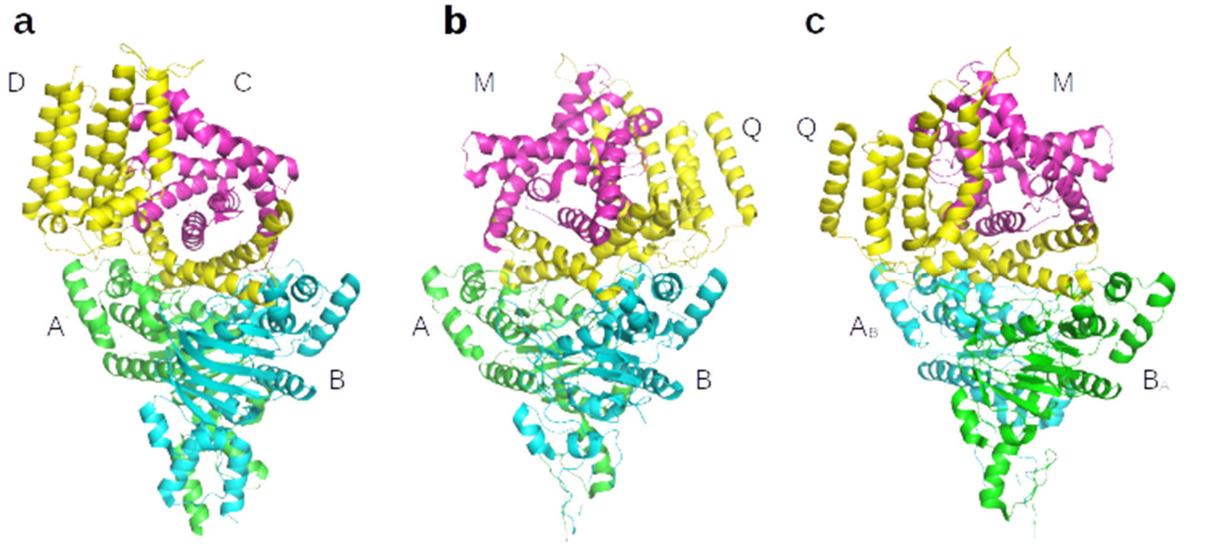


**Supplementary Figure S5: The order of chains influences structural alignment.** If the functional form of the examined protein was built from several chains, then the order and naming of chains in the structure file significantly affected most alignment tools, resulting in wrong structural alignments and scores. For example, when the structures of ECF (Energy Coupling Factor) transporter 7nnu (with chains A, B, C, and D) and 5x41 (with chains A, B, M, and Q) were aligned, the NBDs (named as chains A and B in both structures) overlapped nicely, since they are homologous in nature, therefore it hardly mattered which of the two chains were compared to each other. On the other hand, the two aligned TMDs exhibited a mirrored configuration, since chain D was aligned to chain M and chain C to chain Q (a, b). The same issue occurred with various structural alignment algorithms (PyMOL's align, fit, and super). Simply renaming chain A to chain B and vice versa in 5x41 (reversing the order of NBDs and thus reversing which TMD chain gets aligned to the other structure's corresponding chain) resulted in a correct structural alignment (c). Such chain assignments can be done manually in TM-align by command line arguments. This issue will likely become more significant in the near future, since more structures of complexes are expected to be available thanks to AlphaFold-Multimer. MM-align (https://zhanggroup.org/MM-align) solves this issue by concatenating all chains in all combinations and by performing the alignment in all possible combinations. Since this approach faces serious performance issues, we would encourage the development of tools that automatically match corresponding chains in target structures so that these may be solved unambiguously in the case of many complexes by sequence comparison.

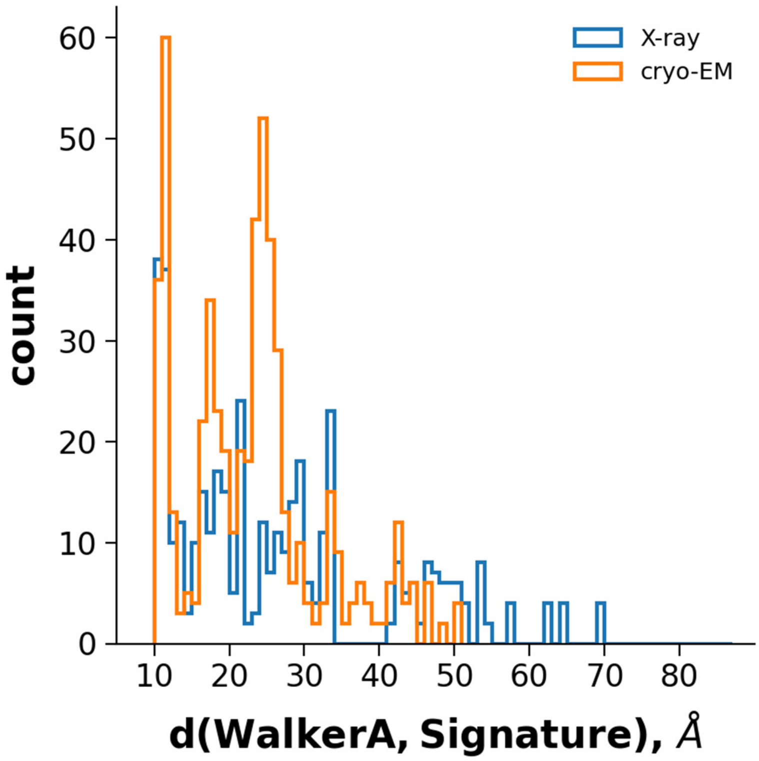


**Supplementary Figure S6: Distances between NBDs grouped by structure determination method.** The distributions of the length of conftor(WA/SIG) were plotted for both X-ray and cryo-EM structures. Distances larger than 51 Å were not observed in cryo-EM structures.

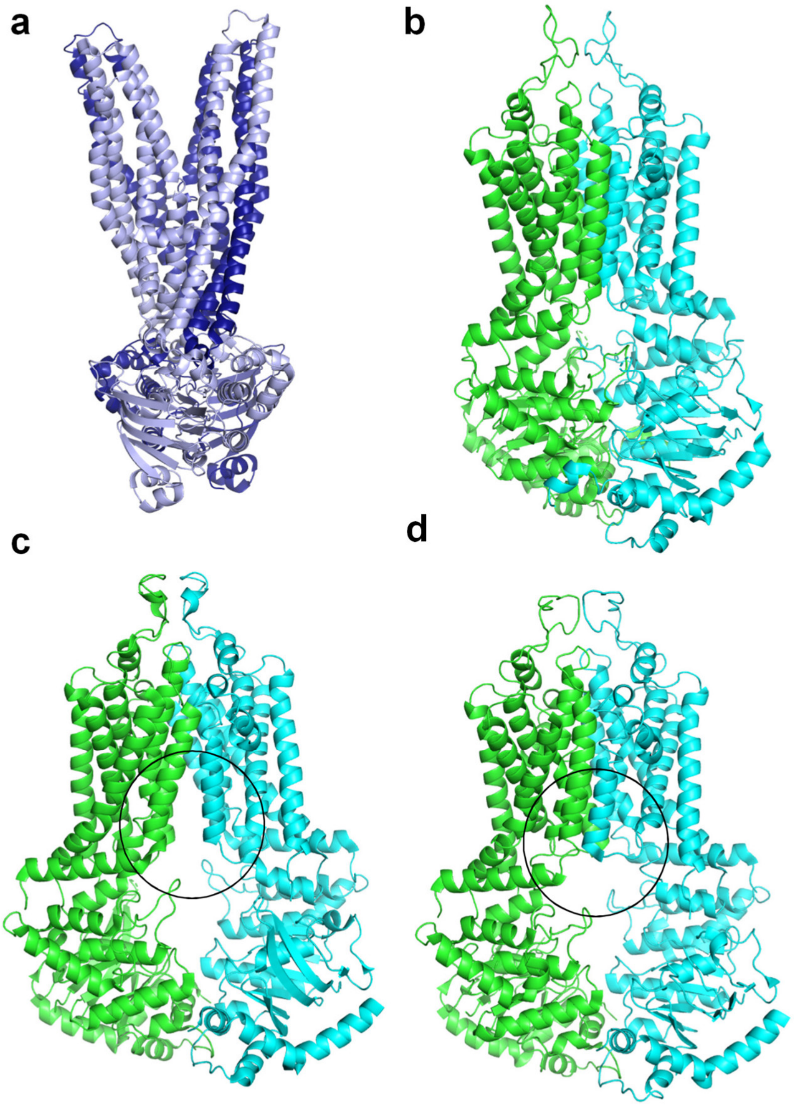


**Supplementary Figure S7: The conformational space is continuous.** (**a**) The ATP-bound Sav1966 structure exhibits a widely open outward-facing conformation. (**b**) ATP-bound ABCG2 structures, such as PDB ID: 6hzm, do not have a large extracellular opening. (**c**) The 6hij inward-facing ABCG2 structure exhibits access to the central drug binding pocket from the intracellular space. (**d**) An inward-facing apo ABCG2 structure (PDB ID: 6vxf) does not have an open gate to the main binding pocket. The main difference between 6hij and 6vxf is circled.

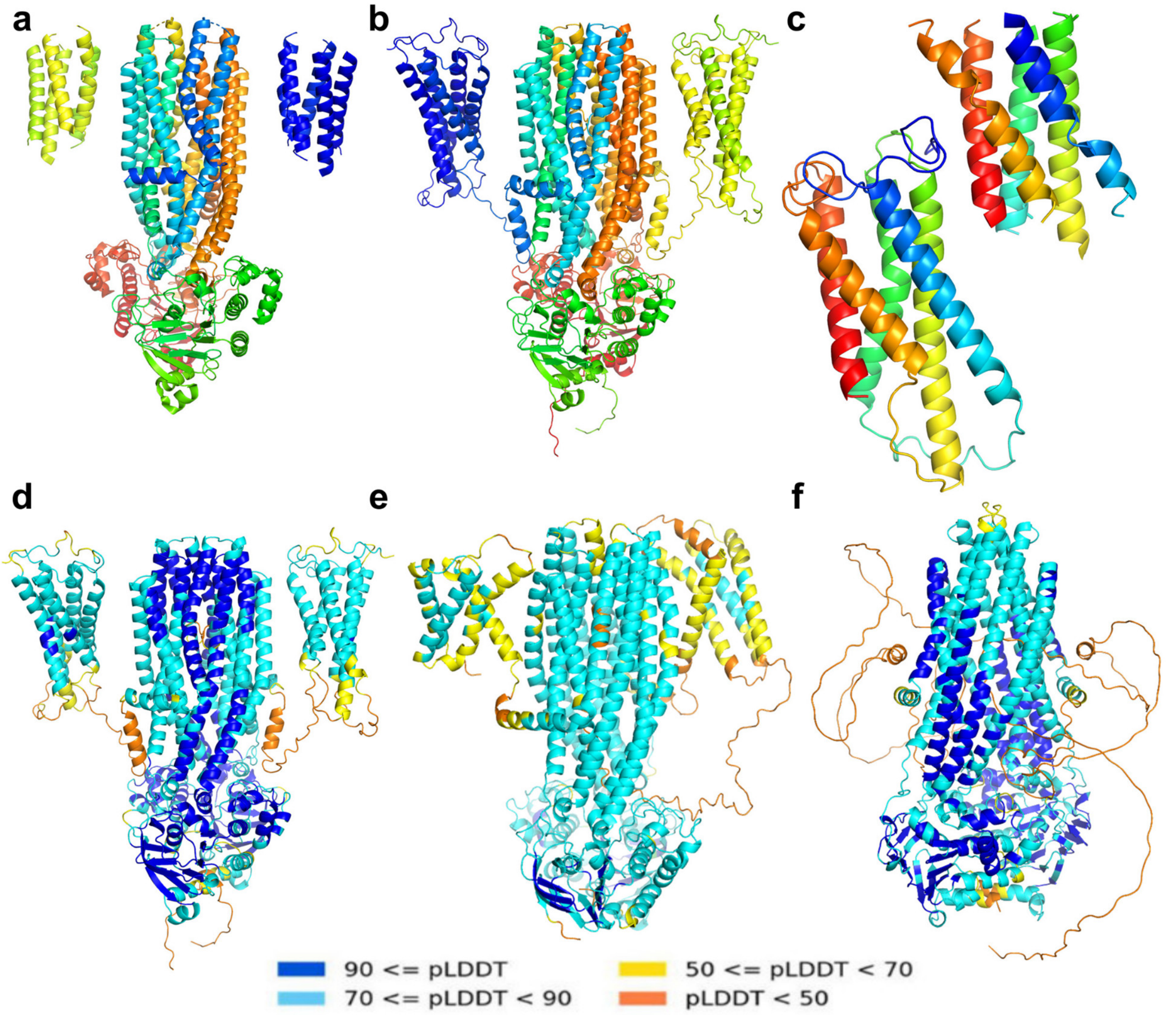


**Supplementary Figure S8: AF2 predictions of ABC proteins.** (**a**) TMD0s (yellow and blue) of ABCB6 are resolved only in a low resolution structure (7d7n, 5.20 Å). (**b**) The AF2-predicted ABCB6 structure highlights the relative positioning of TMD0s. (**c**) The conformation of experimental (top) and AF2 (bottom) TMD0 domains are highly similar, colored from N- to C-termini (blue to red). (**d**) The high pLDDT scores indicate a reliable AF2 prediction. (**e**) In the case of AF2-predicted ABCB2/ABCB3 the pLDDT scores of TMD0 residues are lower, thus the conformation of these domains may be inaccurate and should be handled carefully. (**f**) Among other ABC proteins, ABCB10 also exhibits a long, cytosolic, N-terminal region not resolved in homodimeric experimental structures. Although this region is modeled by AF2, its conformation is wrong, since a significant part of it is unrealistically located in the volume of the transmembrane region. We removed these types of unrealistic regions from ABC structures published via our 3D-Beacon client. The full predictions are also available at http://3dbeacon.hegelab.org. (d-f) Structures are colored according to pLDDT scores.

**Supplementary Table S1: Inward- and outward-facing conformations.**

| fold family | conformational state | reference structure (PDB ID) | n(true) | n(RMSD) |
|---|---|---|---|---|
| Pgp | closed | 6c0v | 66 | 92 |
| | open | 7a69 | 118 | 92 |
| ABCG2 | closed | 6hbu | 8 | 8 |
| | open | 6hij | 37 | 37 |
| MalFG | closed | 3puv | 10 | 10 |
| | open | 3fh6 | 28 | 28 |
| BtuC | closed | 4fi3 | 2 | 3 |
| | open | 2nq2 | 8 | 7 |
| EcfT | closed | 4huq | 6 | 6 |
| | open | 5jsz | 5 | 5 |
| LptFG | closed | 6s8g | 0 | 0 |
| | open | 6s8n | 5 | 5 |
| MacB | closed | 7mdy | 8 | 7 |
| | open | 7mdx | 8 | 9 |
| Bce | closed | 7tcg | 1 | 1 |
| | open | 7tch | 1 | 1 |
| MlaE | closed | 7ch0 | 2 | 2 |
| | open | 7cge | 17 | 17 |

**Supplementary Table S2: AlphaFold-predicted dimers of human half ABC transporters.**

| proteins | structural novelty of the prediction |
|---|---|
| ABCB2-ABCB3 | Two TMD0s are predicted, which are not resolved experimentally (5u1d). |
| ABCB6-ABCB6 | A more complete TMD0 compared to the 7d7n low resolution structure. |
| ABCB7-ABCB7 | Minor compared to 7vgf. Reasonable EL1 is built. The N-terminus (a.a. 1-116) is likely disordered (no TMD0). |
| ABCB8-ABCB8 | Minor compared to 7ehl and 5och. Small unresolved loops are modeled. The N-terminus (a.a. 1-134) is likely disordered (no TMD0). |
| ABCB9-ABCB9 | Two TMD0s are predicted. |
| ABCB10-ABCB10 | Minor compared to 3zdq, 4ayt, 4ayx, and 4ayw. The N-terminus (a.a. 1-150) is likely disordered (no TMD0). |
| ABCD1-ABCD1 | Minor compared to recently determined experimental structures. The N-terminus is falsely predicted to contain an additional TM helix (a.a. 10-38). This segment contains a hydrophobic patch (a.a. 19-27, AVLLALAAY), which is likely immersed into the membrane bilayer or participates in either intramolecular or intermolecular interactions. This region may be a MemMoRF (https://memmorf.hegelab.org; Csizmadia et al. Nucleic Acid Research 2020, gkaa954). |
| ABCD1-ABCD2 | Experiments indicate that ABCD1 and ABCD2 can form a heterodimer. An extra helix traversing the TM region similar to ABCD1 N-terminus was predicted. |
| ABCD2-ABCD2 | An extra helix traversing the TM region similar to ABCD1 N-terminus was predicted. |
| ABCD3-ABCD3 | An extra helix traversing the TM region similar to ABCD1 N-terminus was predicted. |
| ABCD4-ABCD4 | Short N-terminus without extra helix. |
| ABCG1-ABCG1 | Minor compared to recently determined experimental structures (7oz1, 7r8c, and 7r8d). The linker region does not exhibit two helices forming a V-shape, thus it differs from that of ABCG2 and ABCG5/8. |

| | |
|---|---|
| ABCG2-ABCG2 | Minor compared to experimentally determined structures. This prediction demonstrates the quality of the linker region with a double helix V-shape. |
| ABCG4-ABCG4 | The linker region does not exhibit two helices forming a V-shape, thus it differs from that of ABCG2 and ABCG5/8. |
| ABCG5-ABCG8 | Minor compared to recently determined experimental structures. The ABCG5 linker is V-shaped, while the ABCG8 linker is not. |
